# The effect of light vs dark coat color on thermal status in Labrador Retriever dogs

**DOI:** 10.1101/639757

**Authors:** Caitlin Neander, Janice Baker, Kathleen Kelsey, Jean Feugang, Erin Perry

## Abstract

Although dark coat color in dogs has been theorized as a risk factor for thermal stress, there is little evidence in the scientific literature to support that position. We utilized 16 non-conditioned Labradors (8 black and 8 yellow) in a three-phase test to examine effects of coat color on thermal status of the dog. Rectal, gastrointestinal (GI), surface temperature, and respiration rate measured in breaths per minute (bpm), were collected prior to (Baseline — phase 1) and immediately after a controlled 30-minute walk in an open-air environment on a sunny day (Sunlight — phase 2). Follow up measurements were taken 15 minutes after walking (Cool down – phase 3) to determine post-exposure return to baseline. No effect of coat color was measured for rectal, gastrointestinal or surface temperature, or respiration (P > 0.05) in dogs following their 30-minute walk. Temperatures increased similarly across both coat colors (rectal 1.88 °C and 1.83 °C; GI 1.89 °C and 1.94 °C; eye 1.89 °C and 1.94 °C; abdominal 2.93 °C and 2.35 °C) for black and yellow dogs respectively during the sunlight phase (P > 0.05). All temperatures and respiration rates decreased similarly across coat colors for rectal (0.9°C and 1.0°C) and GI (1.5 °C and 1.3°C) for black and yellow dogs respectively (P > 0.05). Similarly, sex did not impact thermal status across rectal, gastrointestinal or surface temperature or respiration rates measured (P > 0.05). These data contradict the commonly held theory that dogs with darker coat color may experience a greater thermal change when exposed to direct sunlight compared to dogs with a lighter coat color.

## Introduction

Darker coat color has been suggested as a potential risk factor for heat injury in dogs in several publications [1–4] However, little evidence is available to support this theory, and a majority of these claims appear in the introduction or discussion sections of publications, or in review articles, with no supporting data. One study in Greyhounds reported higher rectal temperatures in darker colored dogs following exercise but utilized greater numbers of males in the darker coat participant group. These males were significantly larger in size than their female cohorts, so it is not known if sex or size played a role in their results. In addition, the darker colored group (n = 166) had more than twice the number of the light coated group (n = 63) which may have impacted the outcome [5]. In another study using Newfoundland dogs, researchers tested patches of white and black fur exposed to heat lamps. Authors measured the microclimate of the dog’s coat and reported no significant difference in temperature between white and black fur regions on the dogs [6]. However, this study did not examine two separate groups of dogs with single coat colors (i.e. solid black or white).

Work in cattle has demonstrated an impact on thermal status associated with coat color, but this has not been thoroughly investigated in dogs. Increased solar absorption in darker coated cattle has been demonstrated to increase overall heat gain [7]. Darker cattle exposed to direct sunlight had a surface temperature gain of 4.8 °C, while lighter cattle only increased surface temperature by 0.7°C [8]. Additionally, this study reported increased incidence of elevated surface temperature, respiration, sweating, and heat stress signals in darker colored cattle compared to lighter colored cattle.

The risk of thermal injury to dogs is of significant concern to the veterinary community and is considered a common occurrence especially during the summer months. Evidence to validate the ideas surrounding coat color as a risk factor would be helpful in establishing a better understanding of any increased danger facing dark coated dogs. Assessment of risk for heat injury can only be accurately evaluated by studying dogs that incur heat injury in comparison to dogs that do not, whether prospectively or retrospectively. Prospective studies of this nature are inherently difficult to conduct as our current standards of ethics and animal stewardship generally preclude experimentally induced heat injury in dogs. In addition, given the relatively low incidence of naturally occurring heat injury in any given population of dogs, prospective studies relying on naturally occurring cases would require a significant amount of time to complete. Thus, risk of heat injury is primarily based on observations of normal thermoregulatory reactions to safe levels of thermal stress, typically induced by exercise. In this study, we exposed dogs of light and dark coat colors to mild exercise (i.e. loose leash walk) in direct sunlight to assess thermoregulatory reactions and measure various parameters associated with body temperature and thermoregulation. The objective of this research was to identify the impact of coat color on the thermal status of dogs exposed to direct sunlight and to measure the increase in temperature experienced by black dogs as compared to yellow dogs.

## Materials and Methods

### Animals and Diets

Institutional Animal Care and Use approval (protocol#18-022) was received from Southern Illinois University prior to initiation of the study. The study was conducted in mid-June in Carbondale, Illinois with seasonally typical environmental conditions(mean outdoor temperature 29.34±1.76 °C, 84.81±3.17 °F). Non-conditioned Labrador Retrievers (n = 16) from a single kennel and with similar genetics were recruited for participation in this study. “Non-conditioned” was determined as having daily exercise consisting of 4±1 hours of daily group turnout but the absence of a specific conditioning or exercise program. All dogs utilized came from 2 litters to limit for genetic variability. Dogs had a mean age of 2.73±1.86 years, mean weight 26.6±3.32 kg and a mean BCS of 5.5±1.5. Study participants were maintained on a commercial kibble diet (Victor High Energy, Mid America Pet Food Mount Pleasant, Texas) and fed twice daily for 60 days prior to the study. All dogs were up to date on vaccinations (rabies, bordetella, DHLPP) and received a monthly standardized parasite control regimen (Frontline Plus, Merial France) (Interceptor Plus, Elanco, Greenfield, IN). All study participants received a health screening by a licensed veterinarian prior to inclusion in the study and were also assigned a body condition score (BCS) by a trained researcher (Nestle Purina Petcare Company, St. Louis, MO). Following this exam, one canine was excluded from participation due to a previously undiagnosed dermal condition. Dogs of opposite colors were paired according to sex and BCS for participation.

### Phases

The study was separated into three phases. Phase 1 (Baseline) included housing of each dog for 30 uninterrupted minutes in a climate-controlled room in individual crates. Phase 2 (Sunlight) consisted of 30 minutes of loose leash walking at a controlled pace in an uncovered outdoor sandy arena measuring 30m by 60m. The study concluded with Phase 3 (Cooling) and incorporated a 15-minute rest in a climate-controlled room in individual crates. All dogs were monitored throughout the study by veterinary staff stationed in the center of the outdoor arena and climate-controlled holding area, and all dogs were allowed ad libitum access to water while in their crates during both the Baseline and Cooling phases of the study.

Environmental conditions in the outdoor arena and in the climate-controlled room were monitored (Accurite Wireless Weather Station, Chaney Instrument Co. Lake Geneva, WI) to record temperature, humidity and heat index every five minutes.

### Data Collection

Thermal status data for each dog were captured immediately following each of the three phases utilizing four methods as shown in Fig 1. Gastrointestinal (GI) data were captured using an ingestible thermistor orally administered (CorTemp, CorTemp Inc, Palmetto, FL) 30 minutes (±15) prior to the Baseline phase. GI temperatures were monitored with a handheld wireless reader (CorTemp, HQInc Palmetto, FL.). GI temperature was recorded in triplicate for each data collection period to ensure accuracy and the mean was utilized for statistical analysis. Rectal measurements were collected in tandem, using calibrated, 8second digital thermometers (American Diagnostics Company ADTEMP II model #413B) inserted to a depth of approximately 2 cm with petroleum jelly to minimize canine discomfort. Thermal images of participants were captured using a forward-looking infrared thermal camera (FLIR T400 thermal camera) at an approximate distance of 2 meters from the canine to capture body surface temperature as previously described [9–12]. To reduce the effects of environmental factors, all images were captured in an enclosed area with no exposure to wind or direct sunlight. Thermal images were analyzed using thermography software (ThermaCam Researcher Professional 2.9, FLIR Systems Inc. Wilson, OR, USA) to determine body surface temperature at the left eye and caudal abdomen as described previously [13–15] with examples shown in Fig 2.

**Fig 1.**
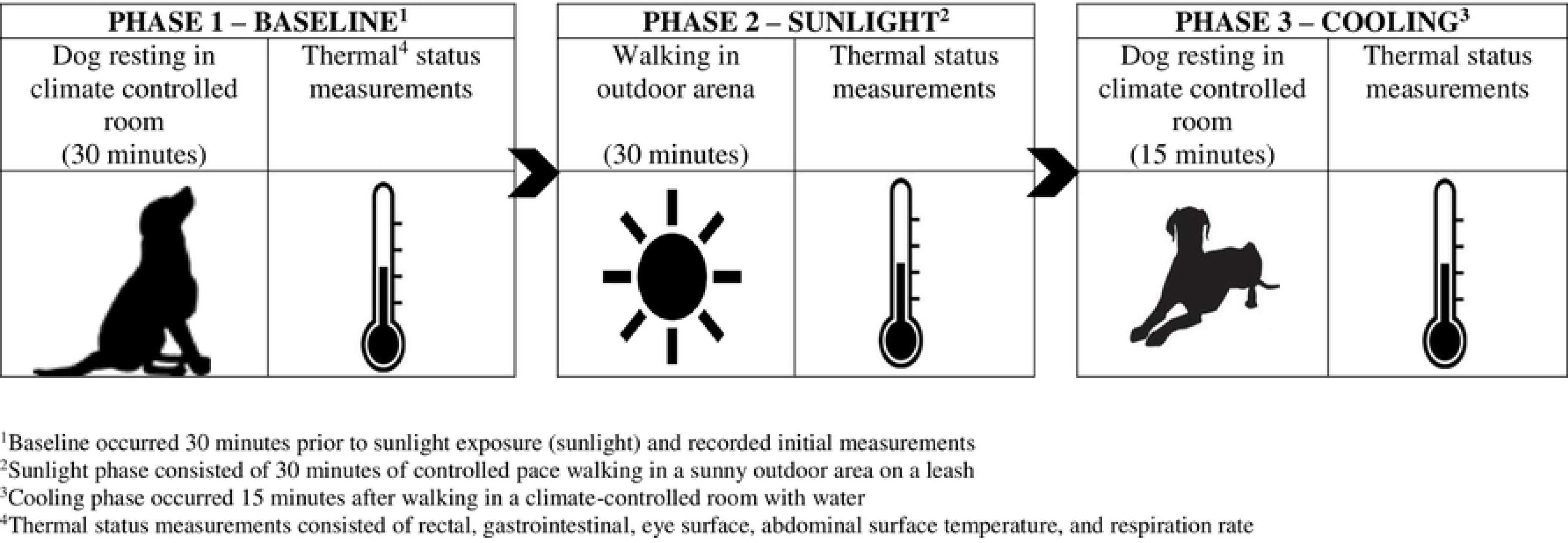
Pictogram representing the phases and points of data collection.

**Fig 2.**
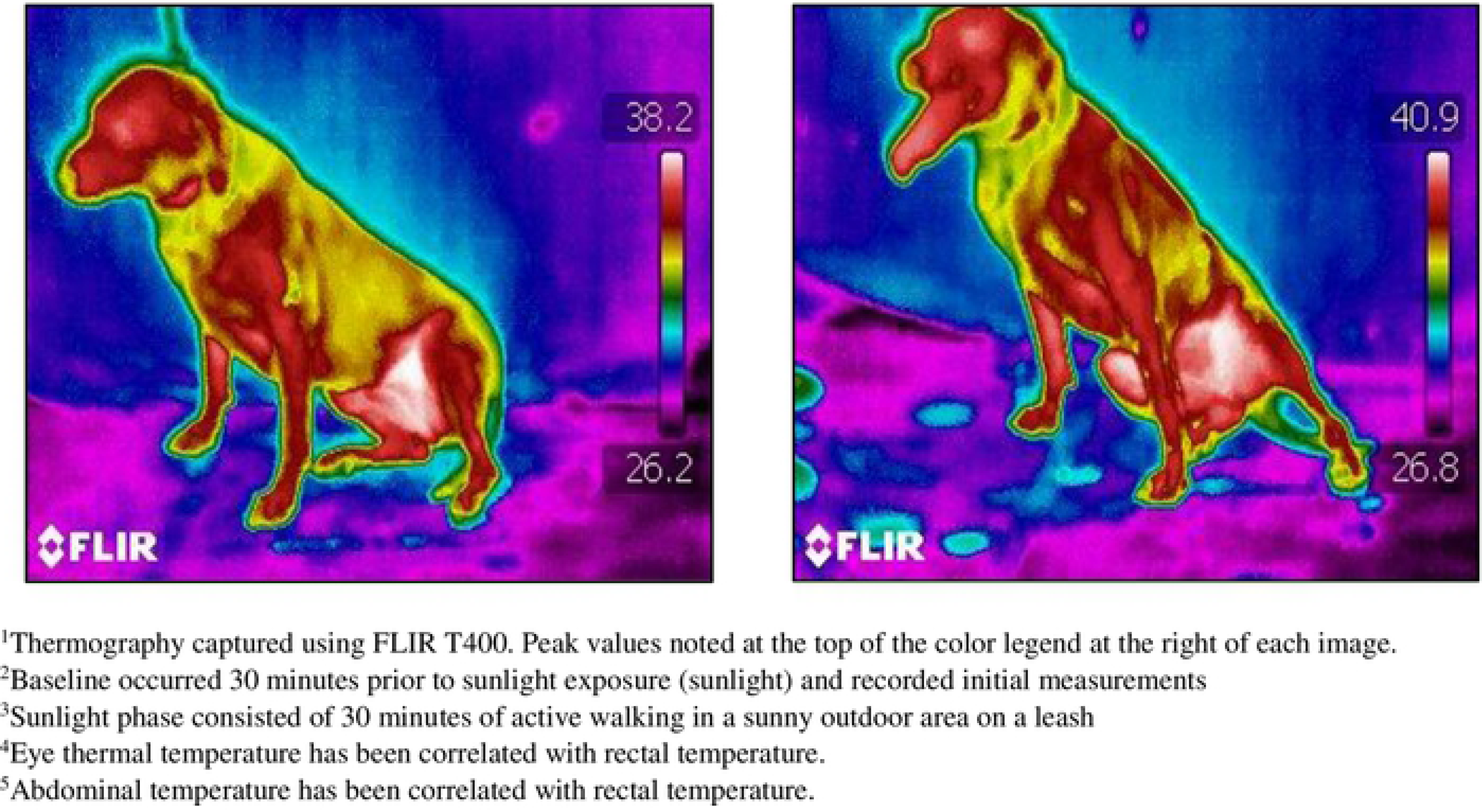
Thermography^1^ depicting Baseline^2^ (left) and Sunlight^3^ (right) values for Labrador Black 6. The two areas of interest were left eye^4^ and left caudal abdomen^5^.

Digital video was utilized to record respiration rates (GoPro Camera, GoPro Inc. San Mateo, Ca) for 30 seconds at the start of each data collection period prior to rectal temperature monitoring. The head, face, and tongue were captured and later played back in slow motion to count respiration during this 30 second period. A single independent observer was utilized throughout all canine respiration videos to minimize observer bias. Respiration was calculated as breaths per minute (BPM) = 30-second respiration × 2.

### Statistical Analysis

All data were analyzed using SAS version 9.4 (SAS Institute Inc., Cary, NC). Each phase (baseline, sunlight, cooling) was examined using a Proc Glm repeated measures test. Baseline and Sunlight temperatures were examined using a paired t test to identify main effects of coat color and sex for dependent variables including rectal, gastrointestinal, surface temperature, respiration rate, and water consumption.

Additionally, a multivariate ANOVA was utilized to identify differences associated with the interactions of coat color and sex on rectal, gastrointestinal, body surface temperature, respiration, and water consumption. Water consumption throughout the data collection period was calculated as: Water offered − Water remaining = Water consumed

Return to baseline was identified as having achieved a cooling phase temperature within 0.5°F of the dog’s initial baseline temperature using the below equations. If the cooling phase temperature had fallen to within 0.5°F, it was deemed “yes” the dog returned to a baseline temperature. The following equation was utilized:

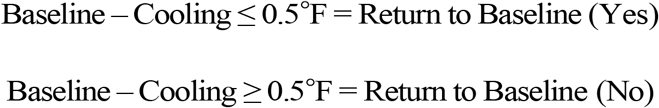

Return to baseline was reported as Yes or No and was analyzed using the Proc Freq procedure of SAS (chi square) to examine differences coat color and sex. Significance for all outcomes was established at P < 0.05.

## Results

Following 30 minutes of walking in direct sunlight, rectal temperatures increased by 1.88°C in black dogs, and 1.83 °C in yellow coated dogs (P < 0.0001). Similarly, GI temperatures increased by 1.89 °C in black coated dogs, and 1.94 °C in the yellow group, (P < 0.0001) as shown in Fig 3. Eye surface temperature increased by 2.8 °C black and 1.93 °C yellow (P < 0.005) and abdominal surface temperature increased by 2.93 °C black and 2.35 °C yellow (P < 0.0001). See Figs 3-4, Table 1. No significant temperature difference was noted between black and yellow Labradors across all phases.

**Fig 3.**
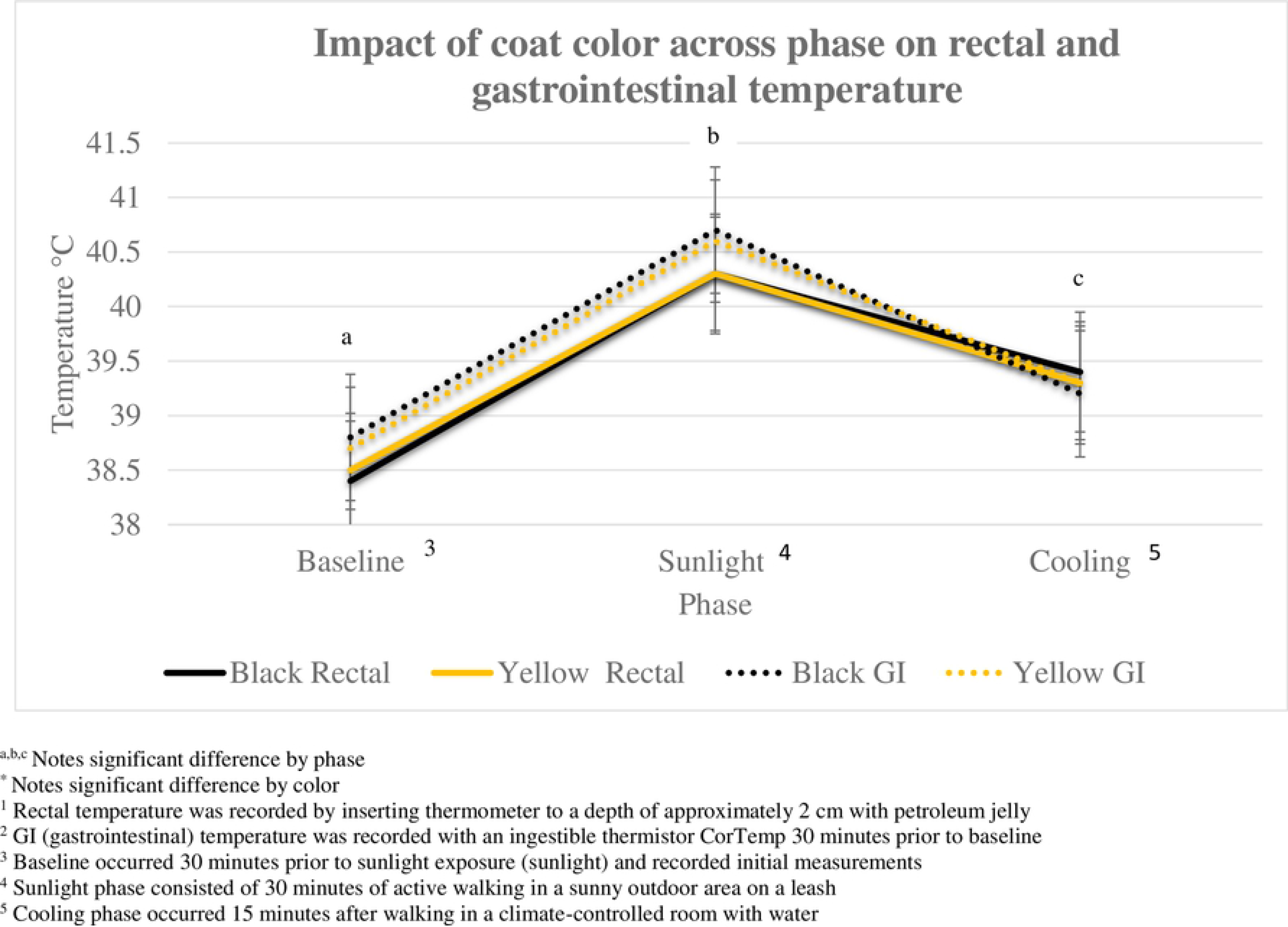
Mean change in rectal^1^ and gastrointestinal temperature (GI)^2^ across three phases in non-conditioned Labradors.

**Fig 4.**
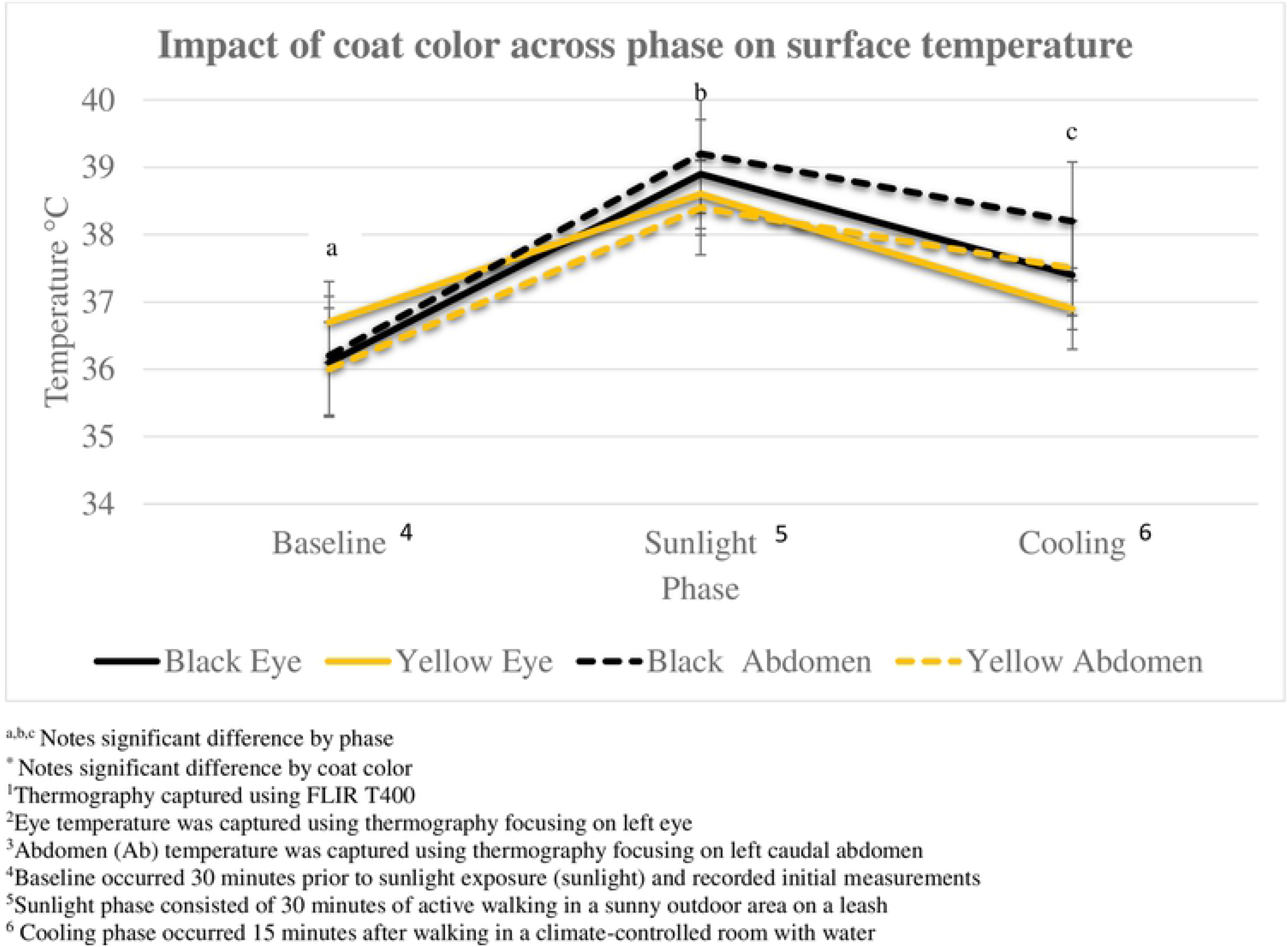
Mean change in body surface temperature measured by thermography^1^ at the eye^2^ and abdomen^3^ in non-conditioned Labradors

**Table 1.** Mean values of thermal status indicators1 across three phases (Baseline^2^, Sunlight^3^, Cooling^4^) in Labradors grouped by coat color.

| Variable | Color | Baseline | P-value | Sunlight | P-value | Cooling | P-value |
| --- | --- | --- | --- | --- | --- | --- | --- |
| **Rectal<sub>5</sub>** | Black | 38.44±0.37 °C<sup>a</sup><br>101.2±0.7 °F | 0.8404 | 40.3±0.41 °C<sup>b</sup><br>104.5±0.8 °F | 0.9354 | 39.46±0.34 °C<sup>c</sup><br>103±0.6 °F | 0.4673 |
|  | Yellow | 38.51±0.52 °C<sup>a</sup><br>101.3±1.0 °F |  | 40.31±0.34 °C<sup>b</sup><br>104.6±0.6 °F |  | 39.43±0.30 °C<sup>c</sup><br>102.8±0.4 °F |  |
| **GI<sub>6</sub>** | Black | 38.76±0.25 °C<sup>a</sup><br>101.8±0.5 °F | 0.6279 | 40.68±0.47 °C<sup>b</sup><br>105.2±0.9 °F | 0.8286 | 39.2±0.6 °C<sup>c</sup><br>102.6±1.1 °F | 0.8153 |
|  | Yellow | 38.66±0.48 °C<sup>a</sup><br>101.6±0.9 °F |  | 40.61±0.21 °C<sup>b</sup><br>105.1±0.4 °F |  | 39.29±0.26 °C<sup>c</sup><br>102.6±0.5 °C<sup>c</sup> |  |
| **Eye<sub>7</sub>** | Black | 36.09±0.57 °C<sup>a</sup><br>97.0±1.1 °F | 0.0934 | 38.89±0.42 °C<sup>b</sup><br>102.0±0.8 °F | 0.3004 | 37.4±0.5 °C<sup>c</sup><br>99.3±0.96 °F | 0.0973 |
|  | Yellow | 36.68±0.55 °C<sup>a</sup><br>98.0±1.1 °F |  | 38.6±0.54 °C<sup>b</sup><br>101.5±1.0 °F |  | 36.9±0.51 °C<sup>c</sup><br>98.4±1.0 °F |  |
| **Abdominal<sub>8</sub>** | Black | 36.24±0.91 °C<sup>a</sup><br>97.2±1.7 °F | 0.7763 | 39.15±0.73 °C<sup>b</sup><br>102.5±1.4 °F | 0.0908 | 37.4±0.5 °C<sup>c</sup><br>100.7±1.3 °F | 0.1002 |
|  | Yellow | 36.01±1.44 °C<sup>a</sup><br>96.9±2.9 °F |  | 38.4±0.75 °C<sup>b</sup><br>101.1±1.5 °F |  | 36.9±0.51 °C<sup>c</sup><br>99.6±1.2 °F |  |
| **Respiration<sub>9</sub>** | Black | 123.3±20.5<sup>a</sup> | 0.7206 | 270.5±40.2<sup>b</sup> | 0.744 | 241.7±24.5<sup>c</sup> | 0.2998 |
|  | Yellow | 132.9±40.7<sup>a</sup> |  | 276.6±28.2<sup>b</sup> |  | 216.4±59.7<sup>c</sup> |  |

\*Notes a significant difference observed by coat color
<sup>a,b,c</sup>Notes a significant difference observed by phase
<sup>1</sup>Thermal status indicators including rectal, gastrointestinal, eye and abdominal temperature
<sup>2</sup>Baseline occurred 30 minutes prior to sunlight exposure (sunlight) and recorded initial measurements
<sup>3</sup>Sunlight phase consisted of 30 minutes of active walking in a sunny outdoor area on a leash
<sup>4</sup>Cooling phase occurred 15 minutes after walking in a climate-controlled room with water
<sup>5</sup>Rectal temperature was recorded by inserting thermometer to a depth of approximately 2 cm with petroleum jelly
<sup>6</sup>GI (gastrointestinal) temperature was recorded with an ingestible thermistor CorTemp 30 minutes prior to baseline
<sup>7</sup>Eye temperature was captured using thermography FLIR T400 at the left eye
<sup>8</sup>Abdominal temperature was captured using thermography FLIR T400 at the left caudal abdomen
<sup>9</sup>Respiration rate was captured for 30 seconds utilizing a GoPro, depicted as breaths per minute (bpm)

Similarly, all temperature measurements significantly decreased from sunlight to cooling phase. Following cessation of cooling phase, rectal temperatures decreased by 0.84°C black and 1.0°C yellow (P < 0.0001) and GI temperatures decreased 1.45 C in black dogs and 1.33 in yellow dogs (P < 0.0001) as shown in Fig 3. Thermal eye surface temperature decreased 1.49 °C in black and 1.71 °C in yellow dogs (P < 0.005), and abdominal surface temperature decreased by 1.0 °C black and 0.85 °C yellow (P < 0.0001) as shown in Fig 4. A similar change in respiration rates was shown across both coat colors of Labradors, meaning that coat color did not significantly impact breathing rates across phases (P > 0.05), as shown in Fig 5.

**Fig 5.**
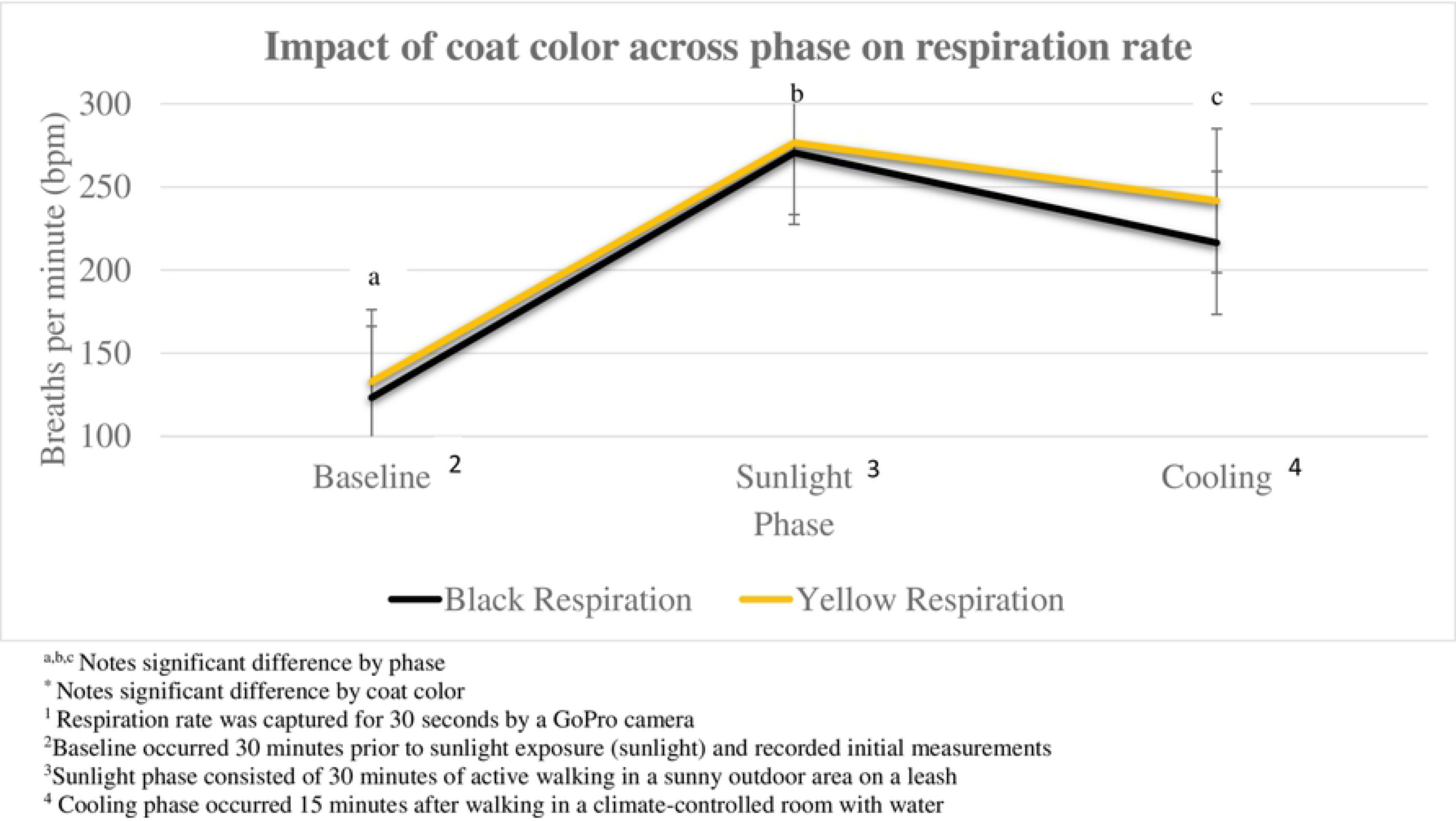
Mean change in respiration rates^1^ across three phases in non-conditioned Labradors.

Exposure to walking in sunlight significantly increased the rectal (1.84°C), GI (1.94°C), eye surface (2.41°C) and abdominal surface (2.67°C) temperatures of all dogs when Baseline and Sunlight temperatures were compared (P < 0.0001). Furthermore, returning to the climate-controlled room significantly decreased the rectal (0.7°C), GI (1.41°C), eye surface (1.58°C), and abdominal surface (0.92°C) temperature of all dogs (P < 0.0001).

After completion of 30 minutes in direct sunlight walking, both males and females saw a similar increase across all temperatures, rectal (1.83°C male, 1.84°C female), GI (2.04°C male, 1.87°C female), eye surface (2.3°C 2 male, 2.5°C female) and abdominal surface (2.93°C male and 2.5°C female). A similar fall in temperatures for both sexes was seen after 15 minutes of passive cooling, rectal (0.92°C male, 0.89°C female), GI (1.32°C male, 1.47°C female), eye surface (1.62°C male, 1.57°C female), and abdominal surface (1.0°C male, 0.87°C female) as shown in Figures 6–7, Table 2.

**Fig 6.**
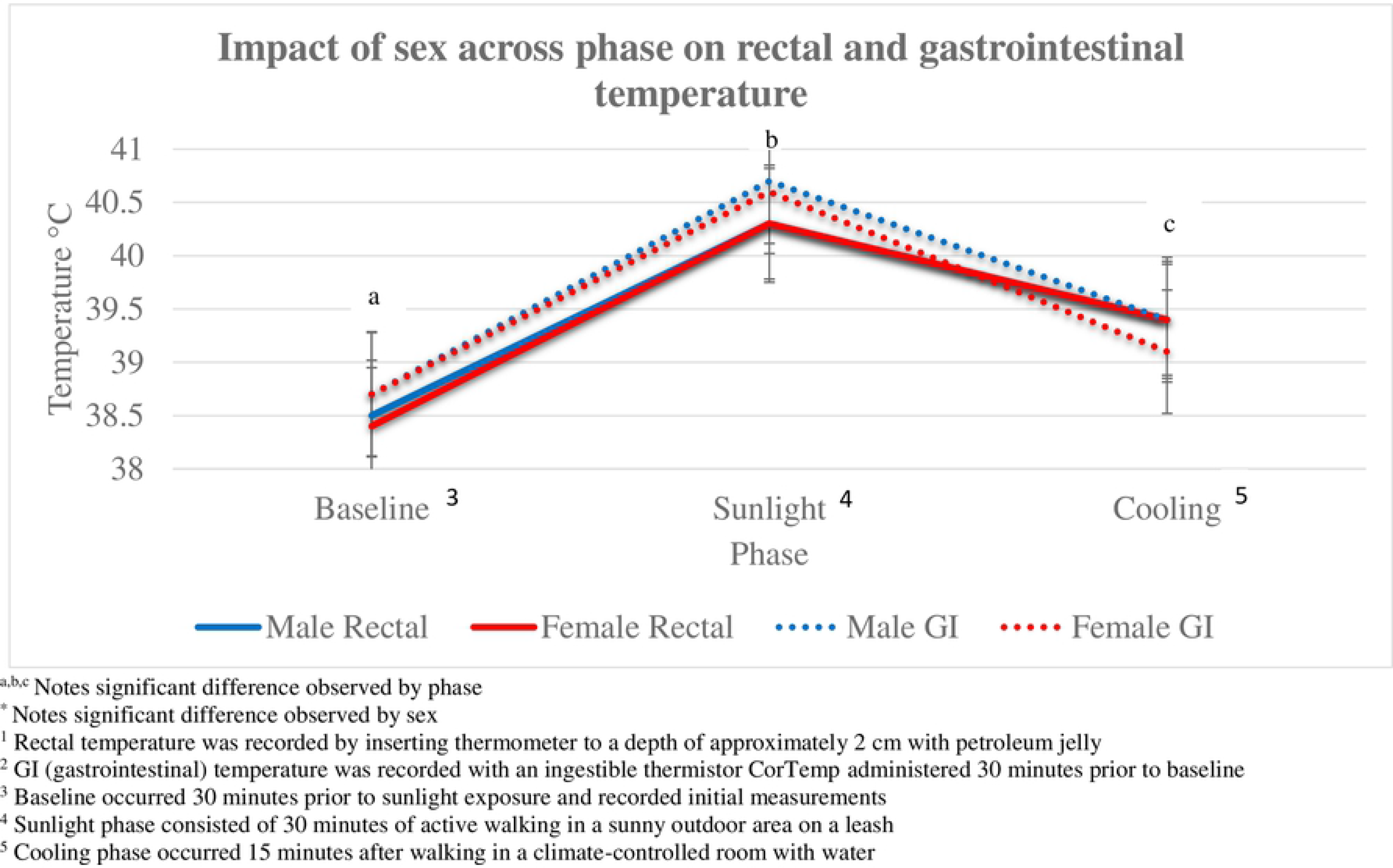
Mean change in rectal^1^ and gastrointestinal temperature (GI)^2^ across three phases in non-conditioned Labradors.

**Fig 7.**
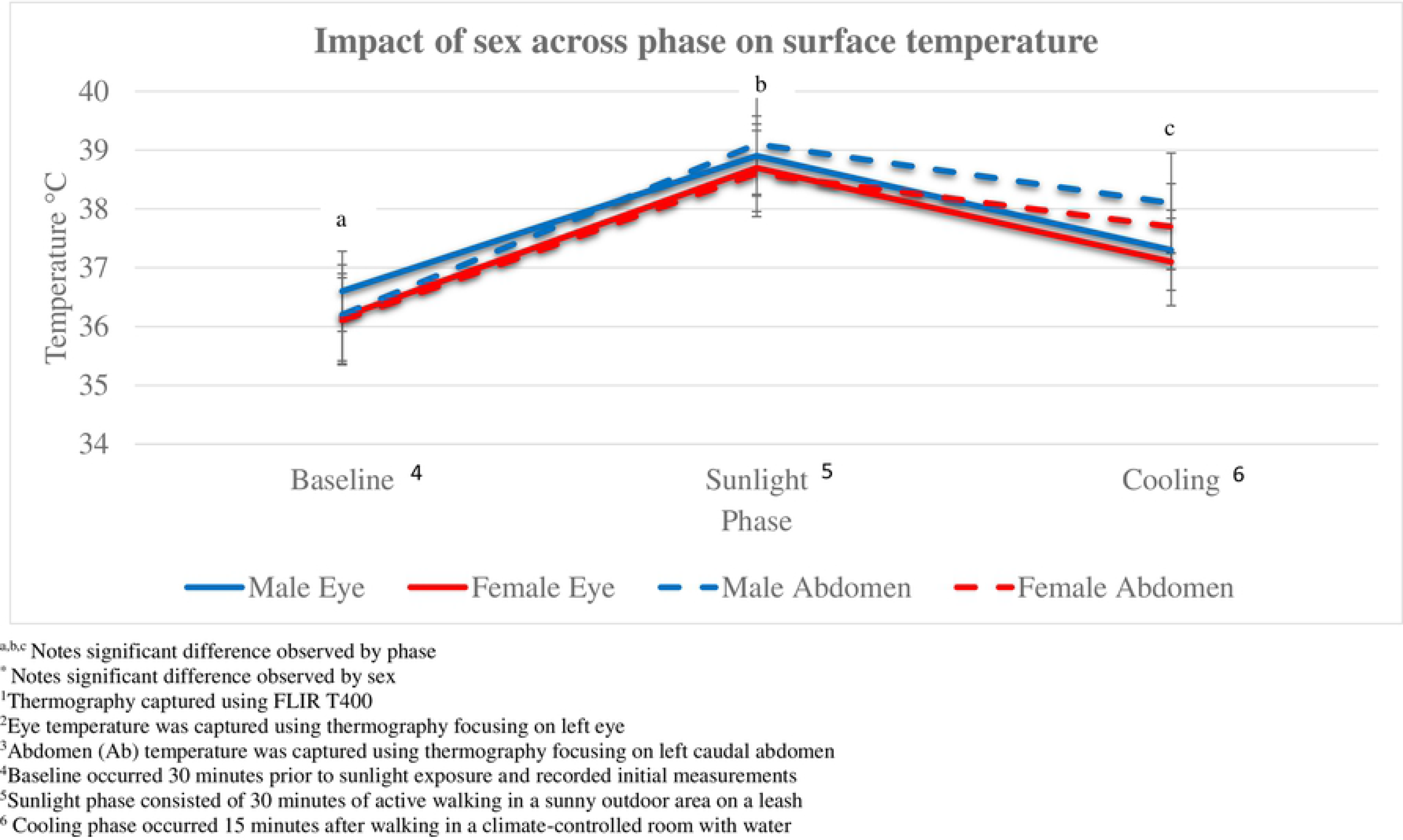
Mean change in surface temperature measured by thermography^1^ at the eye^2^ and abdomen^3^ in non-conditioned Labradors across three phases.

**Table 2.** Mean values of thermal^1^ status indicators across three phases (Baseline^2^, Sunlight^3^, Cooling^4^) in Labradors grouped by sex.

| VARIABLE | SEX | BASELINE | P-VALUE | SUNLIGHT | P-VALUE | COOLING | P-VALUE |
| --- | --- | --- | --- | --- | --- | --- | --- |
| RECTAL<sub>5</sub> | Male | 38.52±0.35 °C<sup>a</sup><br>101.3±0.7°F | 0.8536 | 40.35±0.31 °C<sup>b</sup><br>104.6±0.6 °F | 0.6991 | 39.43±0.3 °C<sup>c</sup><br>103±0.6 °F | 0.7511 |
|  | Female | 38.44±.5 °C<sup>a</sup><br>101.2±0.9 °F |  | 40.28±0.41 °C<sup>b</sup><br>104.5±0.8 °F |  | 39.39±0.27 °C<sup>c</sup><br>102.9±0.5 °F |  |
| GI<sub>6</sub> | Male | 38.68±0.36 °C<sup>a</sup><br>101.7±0.7 °F | 0.9407 | 40.72±0.21 °C<sup>b</sup><br>105.3±0.4 °F | 0.6005 | 39.4±0.41 °C<sup>c</sup><br>102.9±0.8 °F | 0.3736 |
|  | Female | 38.73±0.39 °C<sup>a</sup><br>101.7±0.7 °F |  | 40.6±0.44 °C<sup>b</sup><br>105.1±0.9 °F |  | 39.13±0.49 °C<sup>c</sup><br>102.4±0.9 °F |  |
| **EYE<sub>7</sub>** | Male | 36.58±0.64 °C<sup>a</sup><br>97.9±1.3 °F | 0.2509 | 38.9±0.14 °C<sup>b</sup><br>102.0±0.3 °F | 0.3087 | 37.28±0.5 °C<sup>c</sup><br>99.1±1.0 °F | 0.5457 |
|  | Female | 36.16±0.57 °C<sup>a</sup><br>97.1±1.1 °F |  | 38.66±0.62 °C<sup>b</sup><br>101.6±1.2 °F |  | 37.09±0.59 °C<sup>c</sup><br>98.8±1.1 °F |  |
| **ABDOMINAL<sub>8</sub>** | Male | 36.2±0.95 °C<sup>a</sup><br>97.2±2.0 °F | 0.8360 | 39.13±0.34 °C<sup>b</sup><br>102.4±0.7 °F | 0.1674 | 38.13±0.45 °C<sup>c</sup><br>100.6±0.9 °F | 0.9533 |
|  | Female | 36.09±1.33 °C<sup>a</sup><br>97.0±2.5 °F |  | 38.58±0.97 °C<sup>b</sup><br>101.4±1.9 °F |  | 37.71±0.81 °C<sup>c</sup><br>99.9±1.5 °F |  |
| **RESPIRATION<sub>9</sub>** | Male | 124.0±50.3<sup>a</sup> | 0.8203 | 274.0±34.4<sup>b</sup> | 0.9533 | 225.3±33.9<sup>c</sup> | 0.8553 |
|  | Female | 130.2±51.4<sup>a</sup> |  | 272.9±35.9<sup>b</sup> |  | 230.1±56.0<sup>c</sup> |  |

\*Notes a significant difference observed by sex
<sup>a,b,c</sup> Notes a significant difference observed by phase
<sup>1</sup>Thermal status indicators including rectal, gastrointestinal, eye and abdominal temperature
<sup>2</sup>Baseline occurred 30 minutes prior to sunlight exposure (sunlight) and recorded initial measurements
<sup>3</sup>Sunlight phase consisted of 30 minutes of active walking in a sunny outdoor area on a leash
<sup>4</sup>Cooling phase occurred 15 minutes after walking in a climate-controlled room with water
<sup>5</sup>Rectal temperature was recorded by inserting a thermometer rectally 2cm
<sup>6</sup>GI (gastrointestinal) temperature was recorded with an ingestible thermistor CorTemp 30 minutes prior to baseline
<sup>7</sup>Eye temperature was captured using thermography FLIRT400 at the left eye
<sup>8</sup> Abdominal temperature was captured using thermography FLIR T400 at the left caudal abdomen
<sup>9</sup>Respiration rate was captured for 30seconds utilizing a GoPro, depicted as breaths per minute(bpm)

Across all phases of the study, sex did not show a significant effect on respiration rates of the Labradors, with both sexes showing a similar increase and decrease in bpm (P > 0.05), as shown in Fig 8.

**Fig 8.**
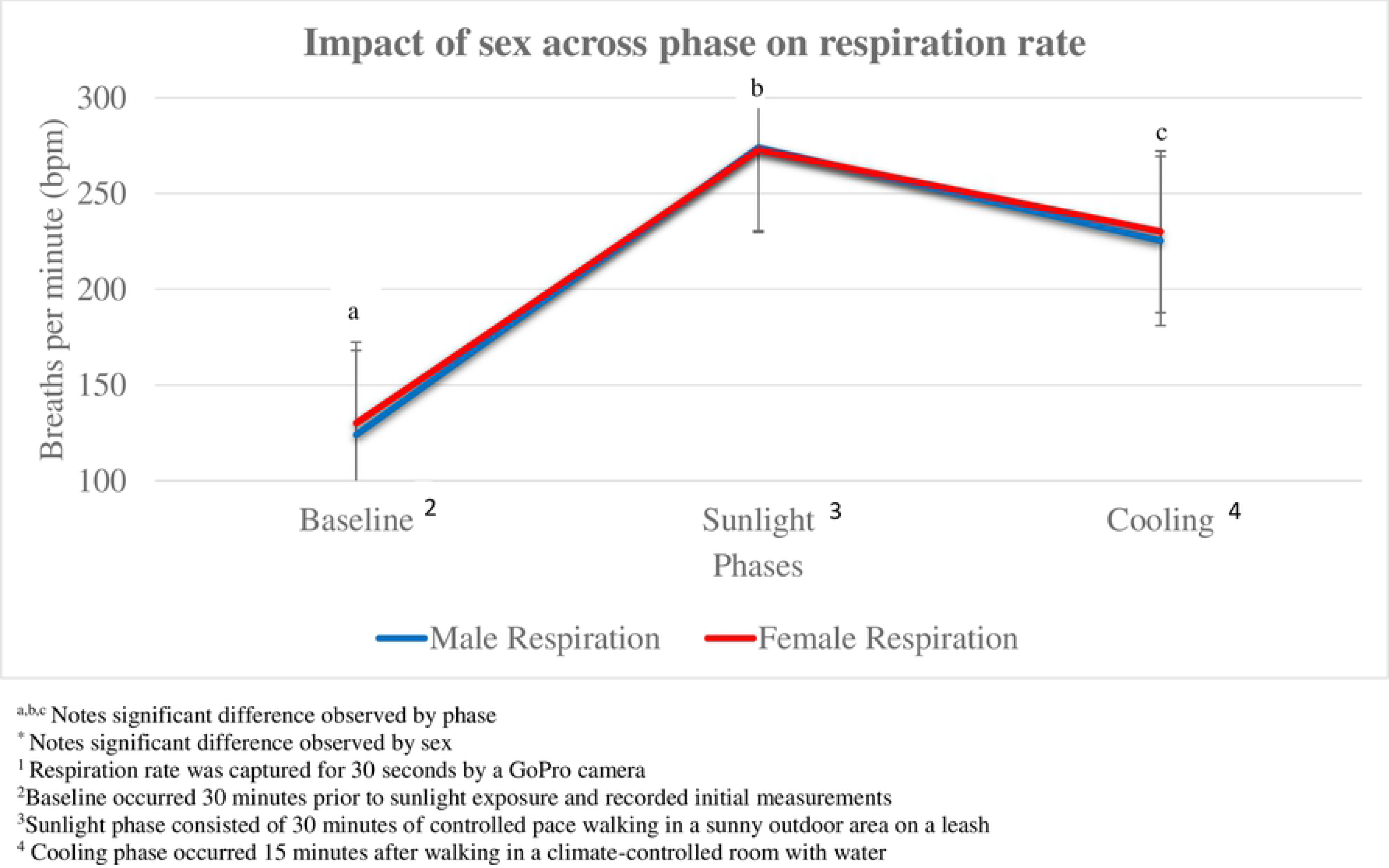
Mean change in respiration rates^1^ across three phases in non-conditioned Labradors.

No effect of coat color (P = 0.5560) or sex (P = 0.9806) was seen for water consumption with black dogs consuming 173.75± 195.3 ml and yellow dogs consuming 221.21±57.1 ml.

No effect of coat color was noted when rectal temperatures were examined for a return to baseline in 12.5% and 28.6% of black and yellow dogs respectively, (P = 0.5692). Similarly, GI temperatures returned to baseline 50% of black and 42.9% of yellow dogs (P = 1.00) and abdominal surface temperature with 12.5% black and 28.6% yellow dogs (P = 0.6080) returning to baseline values. Conversely, coat color did impact the dog’s return to baseline when eye surface temperature was examined with 12.5% black and 71.4% yellow dogs achieving baseline values after their Cooling phase, as shown in Table 3 (P = 0.0406).

**Table 3.** Return to baseline^1^ rectal^2^, gastrointestinal^3^ (GI), thermal eye^4^, and thermal abdominal^5^ temperatures by coat color and sex.

| VARIABLE | BLACK | YELLOW | P-VALUE | MALE | FEMALE | P-VALUE |
| --- | --- | --- | --- | --- | --- | --- |
| RECTAL | 12.5% | 28.6% | 0.5692 | 16.7% | 22.2% | 1.00 |
| GI | 50% | 42.9% | 1.00 | 33.3% | 55.6% | 0.6084 |
| EYE | 12.5% | 71.4% | 0.0406* | 50% | 33.3% | 0.2286 |
| ABDOMINAL | 12.5% | 28.6% | 0.6080 | 0% | 33.3% | 0.6080 |

\*Notes a significant difference between groups
<sup>1</sup>Return to Baseline occurred when cooling temperature returned within 0.5°F of the initial baseline temperature reading measured 30 min prior to sunlight exposure (sunlight
<sup>2</sup>Rectal temperature was recorded by inserting a thermometer in 2cm rectally
<sup>3</sup>GI (gastrointestinal) temperature was recorded with an ingestible thermistor CorTemp 30 minutes prior to baseline
<sup>4</sup>Eye temperature was captured using thermography FLIR T400 at the left eye
<sup>5</sup>Abdominal temperature was captured using thermography FLIR T400 at the left caudal abdomen

Sex did not influence cooling as rectal temperatures returned to baseline in 22.2% and 16.7% of female and male dogs respectively (P =1.00). Temperatures for the GI tract returned to baseline in 55.6% of female and 33.3% of male dogs (P =0.6084).

Similarly, no effect of sex was observed for cooling of surface temperatures measured at the caudal abdomen with 33.3% of female and 0% of male dogs returning to baseline values (P =0.6080). Eye surface temperature returned to baseline in 33.3% female and 50% of male dogs achieving baseline values after their Cooling phase (P = 0.2286), as shown in Table 3.

## Discussion

Black dogs did not demonstrate a difference in temperature following exposure to direct sunlight when compared to yellow dogs for any of the parameters we examined, including rectal thermometer using a standard medical-grade predictive digital thermometer, GI temperature using an ingestible thermistor, eye surface temperature and abdominal surface temperature using forward-looking infrared thermography, respiration or water consumption. Contradictory to currently held beliefs, all dogs experienced a similar rise in rectal, GI, surface temperature, and respiration from the baseline to the sunlight phase, with no difference shown between dark vs lighter coated dogs (P > 0.05) as shown in Figures 3–5 and Table 1. Furthermore, no effect of sex was measured as both males and females demonstrated similar responses to sunlight exposure and cooling based on rectal, GI, eye surface, abdominal temperatures, and respiration rates (P > 0.05) shown in Figures 6-8, and Table 2.

Contrary to the commonly held belief, our data demonstrated that black dogs did not experience a greater heat gain than their yellow counterparts. Similarly, there was no difference in the apparent thermoregulatory effect between dark and light dogs. This is particularly noteworthy because of the relative short duration of the walk and the significant temperature increase we observed in both dark and light-coated dogs. It is also interesting to note that the 15-minute cooling period was inadequate for 80% of the dogs to achieve baseline thermal status based on rectal measurements which are considered standard for recording accurate temperature in animal species [16].

Conversely, almost 50% of each group was able to return to a baseline values via GI values after 15 minutes of cooling. This could be attributed to water consumption during the cooling phase affecting the CorTemp capsule reading. [17] However, it is important to note that all dogs did experience a significant decrease in their rectal, GI, eye and abdominal temperatures, and respiration rates. Future work should include studies with a longer cooling period to determine that time frame necessary for non-conditioned dogs to achieve baseline thermal status following thermal stress. The data presented here are inconsistent with the previous study examining racing greyhounds with larger proportions of dark coated dogs having higher rectal temperatures after racing [5]. Key differences between this study and the prior study on Greyhounds include controlled coat color (black or yellow vs. multiple light or dark colors), tighter grouping of age and sex, and a controlled time period and consistent environment. However, there were fewer dogs in our study compared to the study on Greyhounds (16 vs. 229). Power calculations indicate that it would take more than 500 dogs to adequately test this question using an alpha of 0.05 and 80% power. That number of dogs is beyond our capacity. Furthermore, the greyhounds utilized in the previous study were considerably more fit than the non-conditioned dogs used in our study. Fitness level can impact thermal response as previously demonstrated in working canines [18, 19] and should be examined as a controlled factor in future work.

Infrared thermal cameras have been widely used in livestock species to identify changes in the surface temperature of the animals. These studies focused on areas that had more skin exposure for more accurate data, such as the flank, eye, and facial region. In our canine study, thermal images were captured inside a building to reduce effects from wind, sun, and other environmental exposures. Both yellow and black dogs showed similar changes in body surface temperatures which does not support the idea that coat color is a potential risk factor for thermal stress. A comparison of skin surface temperature during exposure to sunlight in dogs is warranted. There were some challenges associated with the capture of the thermal imaging. Although the baseline photos were captured with little difficulty, many of the dogs were hot following the sunlight exposure and several were non-compliant in assuming the same posture. More obedient/compliant dogs would prove better subjects for this nature of study.

In designing the study, we considered that if body temperature measurements were similar between the dark and light coated dogs, perhaps dark coated dogs simply undertook increased efforts of thermoregulation such as increased respiration (i.e. panting) or increased water consumption. However, we found no difference in these parameters between the dark and light coated dogs, suggesting that the effort they expended to thermoregulation was also similar. A key limitation in our study is that we did not record heart rate, which would be important in assessment of thermoregulatory response. More sophisticated instrumentation and monitoring would be important in further studies to determine if more subtle physiological changes were occurring with thermoregulation.

Novel data produced by this work include an absence of significant difference in body temperature between black or yellow coated dogs. The techniques utilized to assess temperature, panting and water consumption are non-technical, readily available methods for canine handlers or owners to assess thermal status of dogs in the field, and thus are important to prevention of heat related injury. These data provide critical evidence to dispute the theory that dark coat color is a risk factor for thermal stress which is reported across several forums including veterinary textbooks and previously published articles [1–4]. In this experiment, dark and light-colored dogs exposed to the same environment showed a similar heat gain and loss (mean peak rectal 40.31±0.37 °C and mean peak GI temperatures 40.65±0.37 °C respectively). This study also showed that when these non-conditioned dogs reached rectal temperatures near 40.9 °C (105.62 °F), and gastrointestinal temperatures near 42°C (106.16 °F), 15 minutes of rest in a cool room with water available for consumption is adequate to begin to decrease the temperature, although a return to baseline values may not be achieved within that time frame. No medical intervention or active cooling method was needed to decrease the dogs’ temperature and nor heat-related negative health impacts were noted by the veterinary team on site, despite dogs reaching temperatures as high as 42°C (106.16 F).

## Ethical Conflicts

The authors declare that they had no conflict of interest.

## Acknowledgements

The authors would like to thank the undergraduate and graduate students of Southern Illinois University for their participation in this research. The researchers would also like to acknowledge SIT Service Dogs for contributing the use of their dogs for this project.

